# On estimating phenomenological model states for epileptic seizure prediction

**DOI:** 10.1101/2025.03.09.642227

**Authors:** A I Bhatti

## Abstract

The prediction of epileptic seizure, like the disease itself, is a very old but largely unresolved problem. The prediction may highly improve the quality of life for an epileptic patient. A low cost measurement like Electroencephalogram (EEG) involves the non-invasive monitoring of the brain voltage signals to detect the epileptic seizure. This work aims at finding ways to estimate the internal states of the neuron population by looking at the measured EEG signals so that the seizure onset may be predicted in advance. If one may estimate the states of the neural population, then by relating to the bifurcation horizon, one may find the seizure onset time. To find such states one need an estimator/observer of a neuronal state space model. Most of the neuronal models, be it biological or phenomenological, are nonlinear. If a linear or any other approximation is used for the observer design, the bifurcation horizon may not be accurate enough. The biological models of neural population have the barrier of determining all the physiological parameters of a patient which may be bit limiting. Phenomenological neuron model like Epileptor is adapted, which is a nonlinear and discontinuous model, estimating its states may help in finding the bifurcation parameters. However, the nonlinearities are of Lipschitz and monotonic class. Using Linear Matrix Inequality solutions, a Lipschitz Nonlinear model based Observer is developed and tested in simulation, without using approximations of any kind. The simulation shows high fidelity of the observer to the model at hand estimating the states helping in determining the bifurcation parameters.

**Graphical Abstract**

**Highlights:**

- Nonlinear model based observer design for a phenomenological model of epilepsy
- Practical method despite no linear model approximations
- Strong potential for epileptic seizure prediction by estimating the states helpful in making the bifurcation picture.

## 1. Introduction

Epilepsy is one of the oldest known neural diseases in the history of mankind. It can be caused by a variety of malfunctions in the neural circuitry of the brain. It manifests itself in many forms but it is mostly diagnosed by a characteristic signature in the patient’s Electro-Encephalo-Gram (EEG). This signature is in the form of a typical train of pulses repeating at a very low but persistent frequency. In the nonlinear systems parlance, one would be very much tempted to term it as the sign of a limit cycle, though one needs to go deeper to ascertain this conjecture. Human brain, consisting of networks of tens of billions of neurons, is a myriad of webs of nonlinear dynamical systems interacting at several scales. These scales range from a nonlinear dynamical system of a neuronal cell to complex feedback loops of neurons controlling several body functions such as cortico-thalamo-cortical loop (commonly called ctc loop). This ctc loop is responsible for integrating sensory information to the decision making neurons and connecting them back to the neurons controlling muscles. This is just one example from a multitude of feedback loops working on the neuronal network level in the human brain. The mentioned epileptic EEG signals (called eliptoform) may be caused by the gain/phase variations of the components of these feedback loops or even variations in the building block of these networks such as a single neuron. A nonlinear systems scientist may relate to the phenomenon of nonlinear dynamical systems exhibiting characteristic oscillations which are observed at particular instants of time when an otherwise healthy individual undergoes an epileptic episode, commonly called seizures. One is essentially looking at a nonlinear dynamical system expressing markedly different responses including oscillations of specific frequency. It is easy to relate such behavior to the phenomenon of bifurcation i.e. a nonlinear dynamic system showing markedly different behaviors upon the variations in its parameters. During epileptic seizure, one may assume that certain parameters in the neurons in a particular neuronal network loop may undergo a change resulting in the repeating pulse like behavior. The exact dynamics of a seizure are certainly of interest but the likelihood of having a seizure is a fundamental question haunting the medical scientists and clinicians alike for decades. The life would be much easier for the epileptic patients if they know the probability of getting a seizure, while planning for their daily activities or raising the level of precaution. One way to predict seizure may be to devise a pattern of the patients behavior by relating bifurcation analysis to their EEG signals.

These EEG signals are the measurement coming from the neuronal networks of the brain with neurons as their building blocks. If one may observe the states of these neurons or group of neurons by measuring EEG signals, and hence bifurcating patterns, in turn, seizures may be predicted. This takes us to the problem of looking at the state estimators or the observers for the nonlinear dynamic models of neurons. Interestingly, the idea of designing observers for the nonlinear dynamic models of neurons is not new. Such efforts have been going on for many years. More than a decade ago, there was a surge in the efforts of observer design for neuronal models. Not knowing the cause of that trend, such observers could have been used for an attempt at seizure prediction but the literature does not hint in this direction. Before commenting on the nonlinear dynamic aspects of these designed observers, a certain biological fact emerges from the study of these observers. In a classical neuronal model, there are slow dynamics and fast dynamics [1, 2]. The fast dynamics are the Neuronal Output Voltage (coming from Nernst Equation) and a state variable relating to the ion chancel formations. The slow dynamics pertain to the ionic exchanges and distributions both inside and outside of a neuron cell. Most of the neuronal model observers seen in the literature need knowledge of the charge distribution which may be not measured in real time at low cost. Estimation of slow dynamics of a biological model may be a problem. However, the bifurcation analysis performed so far involves both fast and slow dynamics. Thus making a case for designing a neuronal observer estimating both fast and slow states. It would also be of interest to look at the feasibility of such an observer from nonlinear observability standpoint. So far we have motivated the need for a neuronal observer which may estimate and track a neuron cell and predict its states in real time to predict an epileptic seizure. It has been stressed in the literature that predicting slower dynamics is crucial as they represent ionic exchange which is the major cause or precursor to the triggering of fast dynamics of electric pulse train generation which is the hallmark of a seizure. However, most of the neuronal observers designed so far, either focus on the faster dynamics only or do not represent biological model because of the problem of the unknown biological parameters. For this reason, many of the designed observers are built on the basis of phenomenological models. Epileptor is such a phenomenological model which shows high fidelity to EEG data and also is amenable to nonlinear analysis[3],[4],[5]. A vast body of work exists on observer design for nonlinear systems but we will only review the works related to the observer design for neuronal models. There are two main approaches, Kalman Filter based and the other one using geometric control theory. In the Kalman Filter domain all the varieties of Kalman Filter like Extended Kalman Filter (EKF) and Unscented Kalman Filter (UKF) were proposed [6]. This design method has two main aspects which should be carefully analyzed. The model used by the Kalman Filter and its variant are essentially linearized version of the neuron nonlinear dynamics around a particular equilibrium point and if the equilibrium point shifts, the model will essentially change too, thus not helping in any seizure prediction based on bifurcation analysis. In addition, the underlying stochastic distributions are mostly assumed to be Gaussian or be known at the least which is hardly the case. In the case of a neuron, both process and measurement uncertainty are not known apriori. The second approach looks ideal and promising.

In general control theory, a large body of work for nonlinear observer design consists of Nonlinear Lipschitz Observers where a significant part of the model is linear while the nonlinear part comprises of Lipschitz Nonlinearity. If such a method is attempted on a Neuron model then the linear part on its own does not remain observable as the author attempted it for the unified neuron model proposed by depannenmacher [1], [7] and background given in [8], [9]. The nonlinear observer design using geometric control theory was presented by Fairhust and colleagues [10]. They proposed such observers by using second order phenomenological model of Neuron called Rose and Hindmarsh model [11]. A formal framework for such design was presented by Astolfi [12]. Most of the geometric control theoretic approaches are centered on finding a diffeomorphism or nonlinear transformation which may bring the neuronal model into a canonical nonlinear observer form. As Astolfi mentioned himself that this step is not straight forward and may represent a major hurdle in the current pursuit. The onus is on the design of a nonlinear observer utilizing the biological model of the neuron and should be capable of observing and hence estimating all the dynamics of the neuron both faster (electrical) and slower ones (ionic flows). A novel breakthrough is required in the geometric theory of observer design. Another issue is nonlinear observability. Please see the works of Hermann [13], Isidori [14], and others[15], [16] for further details. Regarding biological models a good starting point is the work of Huxley and Hodgkin (H-H model) [17] and a recent addition could be the work by the team of V. Jirsa[1]. It may be added that the task at hand is daunting but feasible. Sotomayor [18] proposed an Extended Kalman Filter (EKF) to be used as an observer for Epileptor which is a phenomenological model of a neuron. Since an EKF depends on a linearized model at every update, it is an approximation to the actual nonlinear model at the best. An observer is needed which may utilize the nonlinear functions and structure of a neuron model, in this case, an Epileptor model. Brogin and colleagues [19, 20, 21] proposed an Epileptor observer by approximating the Epileptor model nonlinearities with a bank of simplified Fuzzy membership functions. The approximation uses differential mean value theorem which may only be used for continuous functions.

There is a clear need for an observer design technique which may take into account the Epileptor nonlinear model per se, without any approximations. As the Epileptor system is discontinuous, nonlinear and exhibits bifurcation. There is a wide body of work dealing with the design of the observers for the systems with Lipschitz or monotonic nonlinearities. These Linear matrix inequalities (LMI) based Observers, despite being linear in structure, can estimate the states of a system with Lipschitz nonlinearities without any approximation. In the past, for such nonlinear systems, the observers may be constructed using the Riccati Equation solution and later on by a convex optimization framework. Hedrick [22] and Rajamani [23] proposed linear observers for the systems having Lipschitz nonlinearities by posing the problem as a Riccati Equation. This was further improved upon by Phanomcheong and Rajamani in 2010 [24]. The problem with the Riccati solution is of finding the right parameters so that the posed Riccati Equation is solvable or feasible. This problem was solved by the LMI-based convex optimization by Boyd [25] and the relevant software yalmip [26] to a larger extent. Arcak and researchers [27] posed an LMI observer for such systems. Arcak devised a method so that the structure of the model nonlinearity becomes a part of the observer thus rendering the observer nonlinear. Zemouche et al used this concept in their LMI-based observer for the systems with Lipschitz nonlinearities and later extended to cover the systems with monotonic nonlinearities [28]. They proposed a way to add further parameterization in the nonlinearity structure so that the LMI problem can be solved more effectively.

This work addresses the design and simulation of an observer of a phenomenological model of an epileptic seizure using an LMI-based observer design for systems with Lipschitz nonlinearity. Section two discusses the design methodology and the results are discussed in Section three. Conclusions are drawn in the last section.

## 2. Methods

The section pertains to the description of the Epileptor model and the subsequent design of its observer based on the Lipschitz nonlinear observer theory. The framework is symbolized if Figure 3

**Figure 1.**
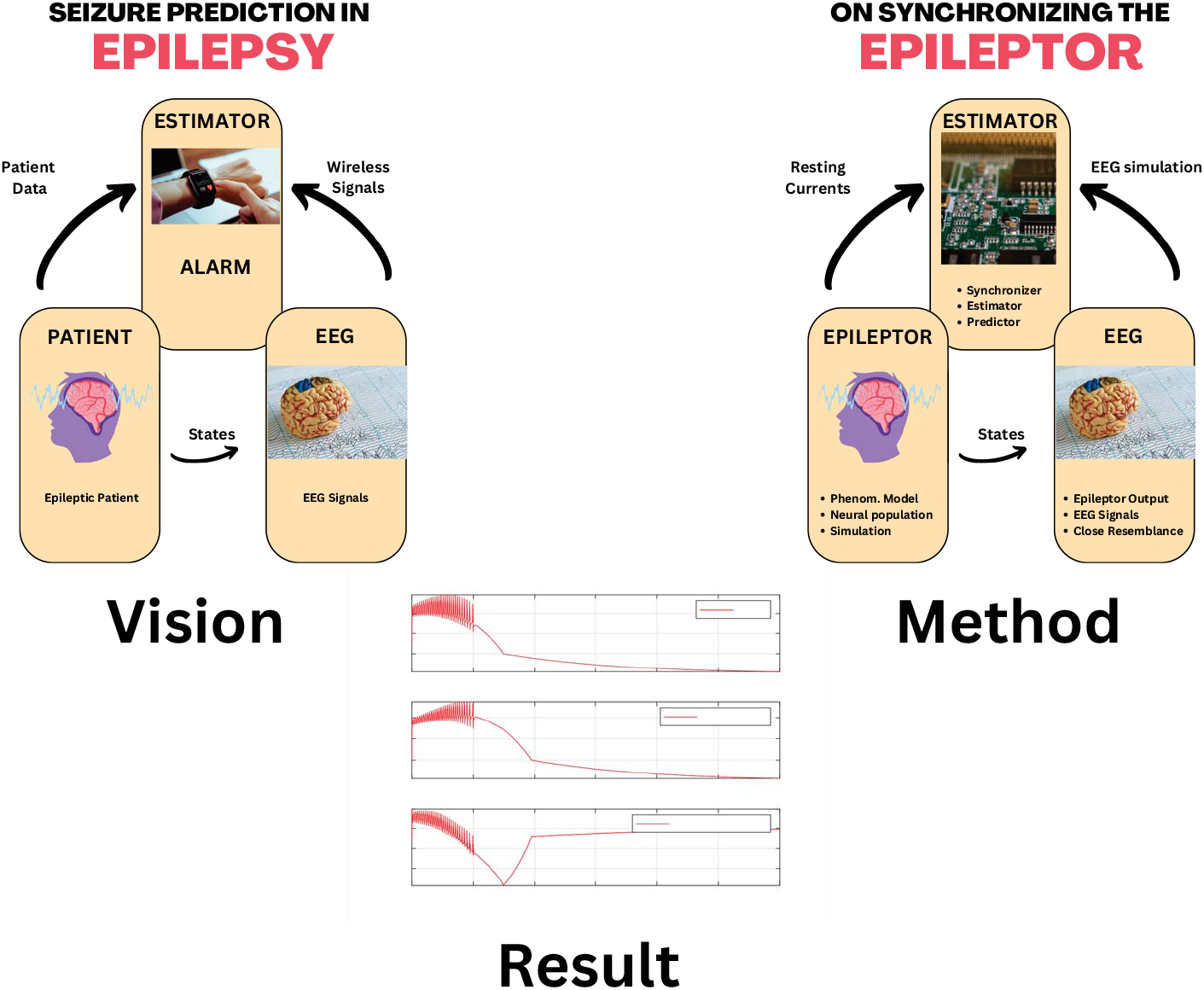
Graphical Abstract

**Figure 2.**
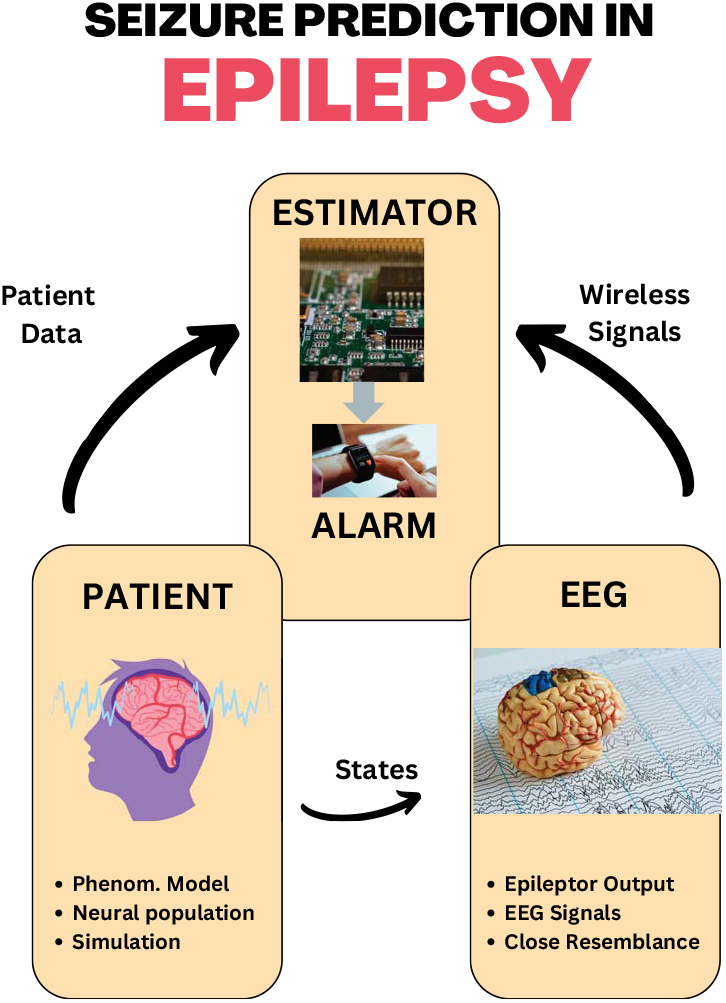
Vision

**Figure 3.**
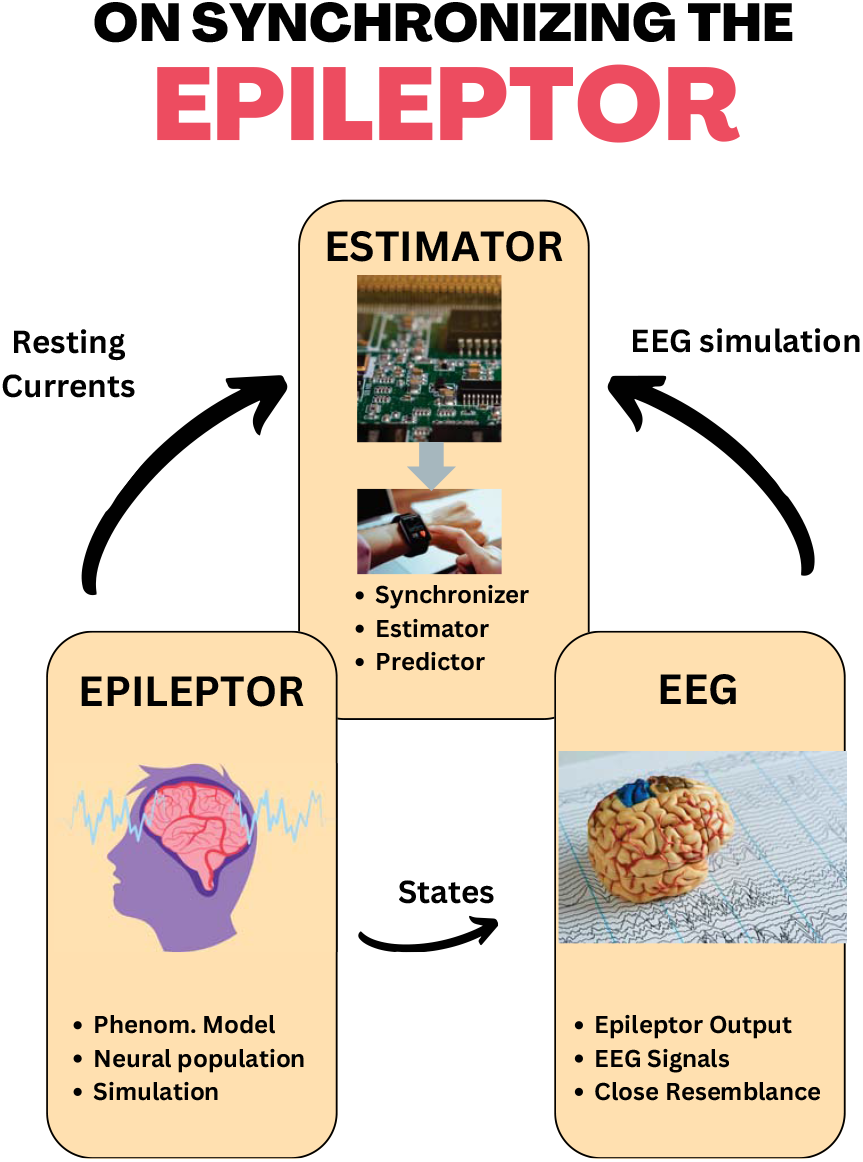
Method

### 2.1. Epileptor Model

The Epileptor model consists of the following set of differential equations:

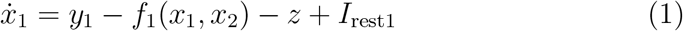

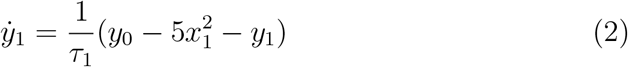

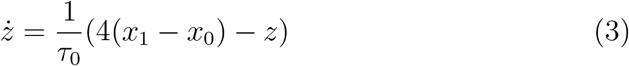

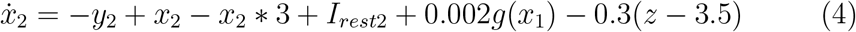

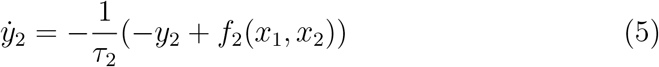

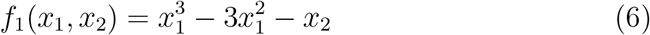

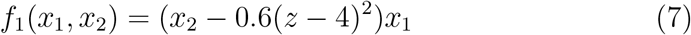

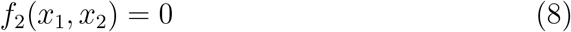

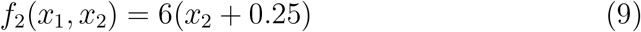

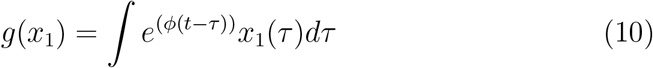

and the parameters are:

- *x*_0_: resting state for *x*_1_
- *y*_0_: resting state for *y*_1_
- *τ*_0_: time constant for *z*
- *τ*_1_: time constant for *y*_1_
- *τ*_2_: time constant for *y*_2_
- *I*_rest1_: resting current for *x*_1_
- ^*I*^_rest2_:

The nonlinear functions in the epileptor model are discontinuous in the model state space. As these discontinuities are on *x*_1_ = 0 and *x*_2_ = *−* 0.25. So these thresholds mark the boundaries dividing the state space into four quadrants, each quadrant following its own sub-model. The four quadrants have these four sub-models which are explained below:

- Quadrant I, *x*1 >= 0, *x*2 >= *−*0.25
- Quadrant II, *x*1 < 0, *x*2 >= *−*0.25
- Quadrant III, *x*1 < 0, *x*2 *< −*0.25
- Quadrant IV, *x*1 >= 0, *x*2 *< −*0.25

The detailed respective sub-models are given in the appendix. The sub-model of Quadrant I is given as:

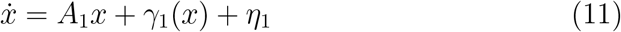

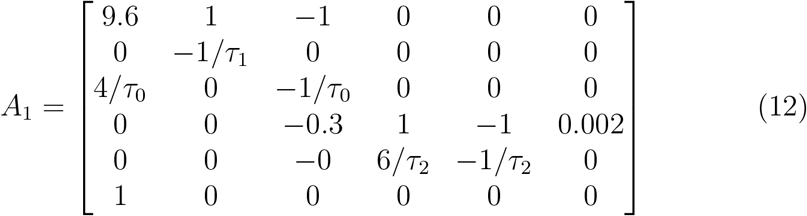

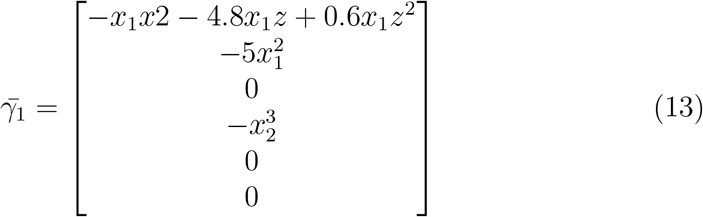

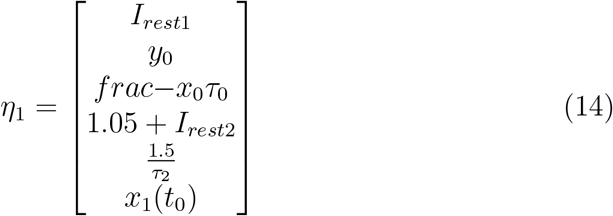

For Quadrant II:

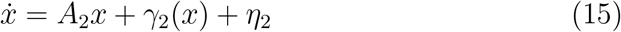

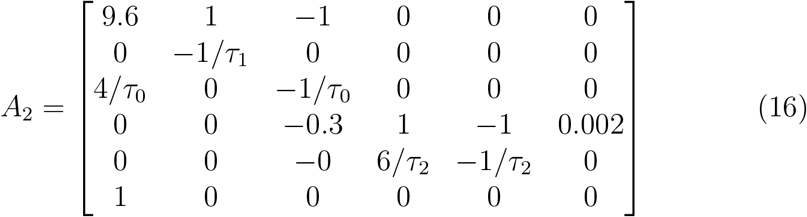

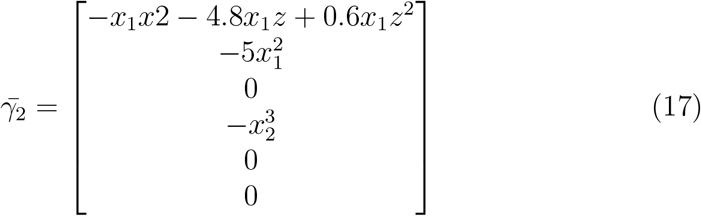

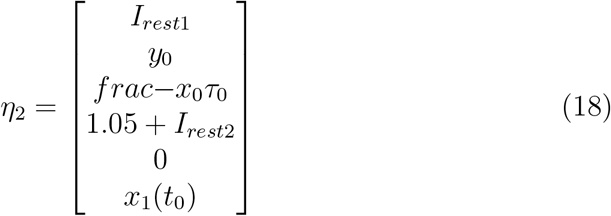

For Quadrant III:

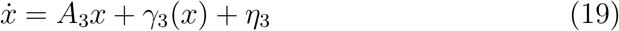

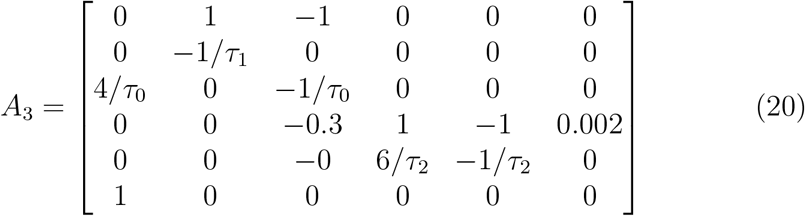

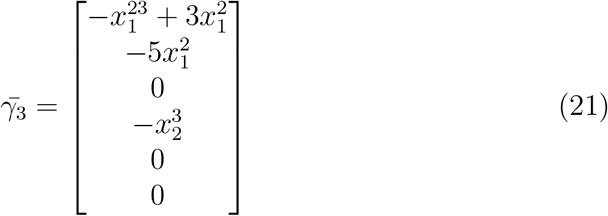

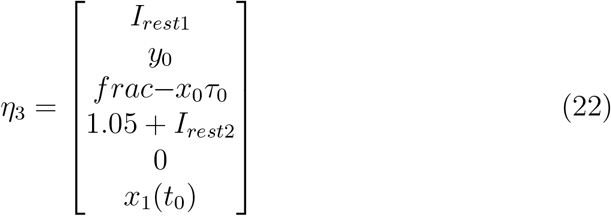

For Quadrant IV:

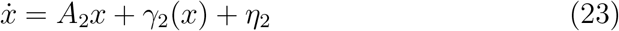

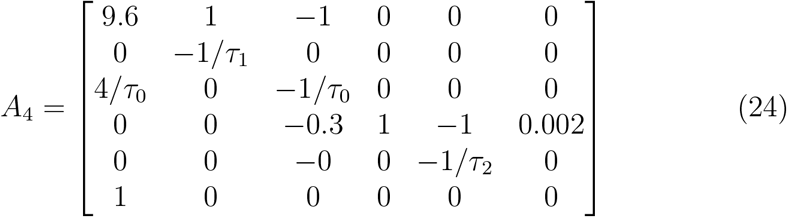

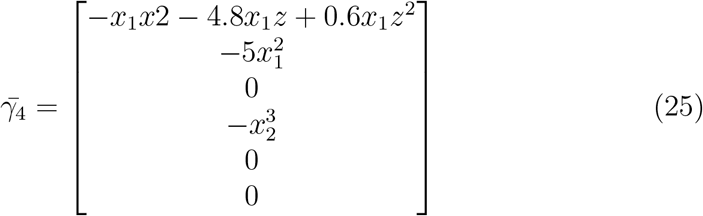

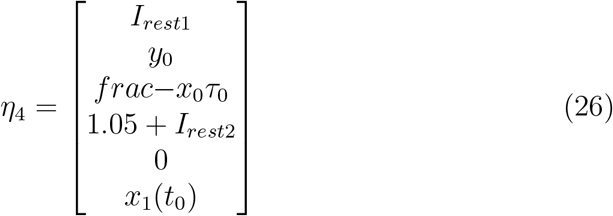

Where *A*_1_ is the system matrix prevalent in Quadrant I, *γ*_1_ is the state nonlinear function relevant to Quadrant I and *η*_1_ is the vector containing external signals and biases. Regarding the states *x*_1_ and *x*_2_ are coming directly from the EEG measurement with your units in mill-volts or microvolts. The output *y* is defined as:

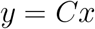

where *C* = [*−*1 0 0 1 0]. Time lags or constants are incorporated through the states *y*_1_, *y*_2_, and *z*. So these time constants express themselves in the pivotal or diagonal elements of the system matrix *A*_1_. These constants also find their way in the bias term *η*_1_. The second set of constants are external signals *I*_*rest*1_ and *I*_*rest*2_. These are important to mention as the nominal values are suggested in [3] but they may change considerably in the actual situations. Similarly, the equilibrium or setpoints *x*_0_ and *y*_0_ may also change. For the LMI-based observer design, one may assume the Linear Parameter Varying (LPV) approach where vertices of the state space of interest are identified and for each of the vertex-model an observer is designed. For the observer implementation, all of the designed observers are then incorporated in an LPV manner. To make the resulting observer easy to implement in real-time, only one observer is designed based on the worst-case scenario. Keeping this in mind, Quadrant I seems appropriate. As per the above discussion the LMI observer for the Lipschitz nonlinear system of Quadrant I is designed in the next Section.

### 2.2. Observer design for Epileptor

The Observer for the epileptor is designed using the methodology proposed by Zemouche and the co-authors [28, 29, 30]. If the system is given as:

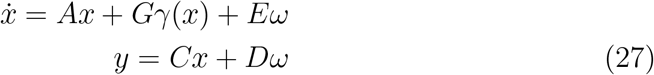

where *n* is the number of states, *m* is the number of nonlinear functions such that : *x ϵ R*^*n*^ are the epileptor states, *γ ϵ R*^*m*^ is the vector of the nonlinear functions *γ*_1_ *γ*_2_..*γ*_*m*_ in the epileptor model,

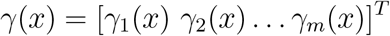

and *ω ϵ R*^*q*^ is the disturbance. The matrix *A ϵ R*^*n×n*^ is the system matrix, *G ϵ R*^*n×m*^ is the distribution matrix for the nonlinear functions, and *E ϵ R*^*n×q*^ is the disturbance distribution matrix. Both *D* and *E* are taken null as the disturbance is assumed to be absent. In the case of the epileptor model it is to be noted that out of the six epileptor states, only few of them are used by each nonlinear function *γ*_*i*_(*x*). From the epileptor model of Quadrant I (Equation 12) the nonlinearities are :

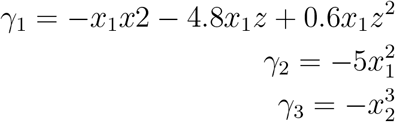

Rewriting the nonlinear functions *γ*_*i*_(*x*) as *γ*_*i*_(*H*_*i*_*x*).

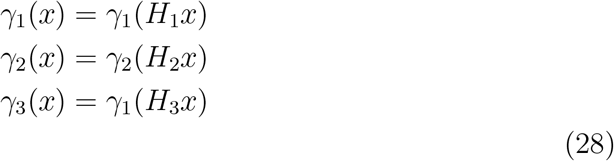

Where

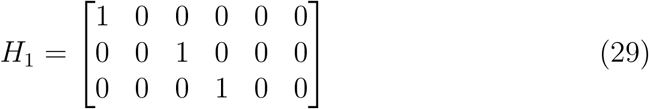

if *n*_1_ is the number of state variables used in *γ*_1_(*x*), then *H*_1_ *ϵ R*^*n*1*×n*^. Similarly:

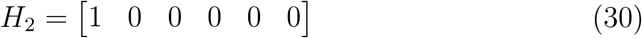

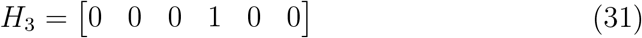

The nonlinearity function distribution matrix *G* for the Epileptor Quadrant 1 model is formulated as:

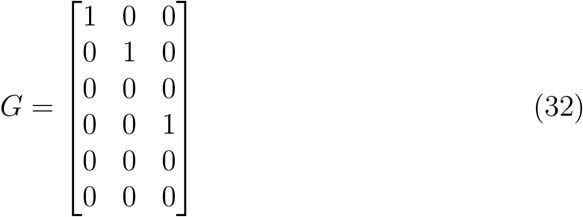

First state is acted upon by *γ*_1_(*x*), the second state by *γ*_2_(*x*) and the fourth state by *γ*_3_(*x*). In the last, the application of Lipschitz constants is facilitated by the matrix 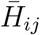 :

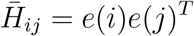

where *e*(*i*) *ϵ R*^*n*^ is a vector with all elements to be zero except for the *ith* entry, which is made equal to 1. This is defined in Lemma 7 of [30]. The proposed observer structure is:

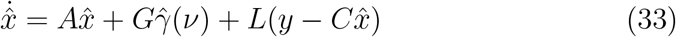

where:

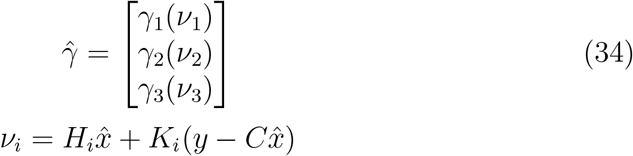

The gain matrices to be designed are *L, K*_1_, *K*_2_, *K*_3_. These are designed by solving the following LMIs:

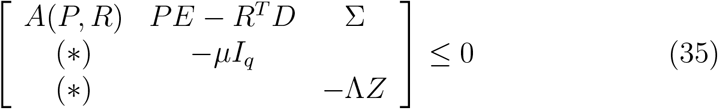

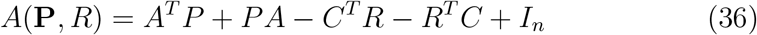

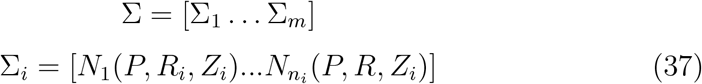

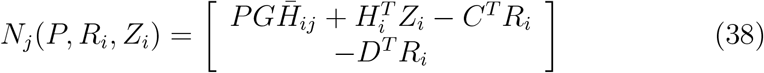

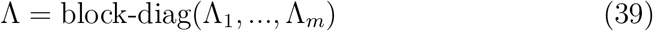

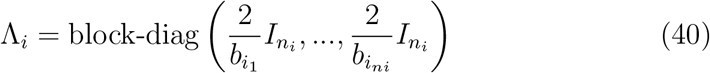

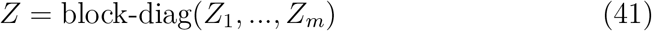

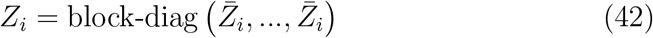

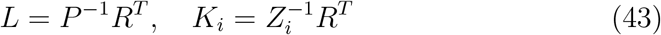

The next section discusses the issues of parameter tuning and observer convergence.

## 3. Results and Discussion

From Figures 4, 5, and 6 it can be seen that the estimates of states *x*_1_ and *x*_2_ converge within ten minutes and the third state *z* takes more time to converge, even in the excess of an hour. This excess time can be attributed largely to the two factors. The first is the large mismatch in the initial conditions of the epileptor and that of its observer which amounts to nearly twenty percent of the final value of *z*. This mismatch is deliberately introduced to demonstrate its damaging effect. It can be assumed that in lab or clinic conditions this observer will be allowed to run for a while under the healthy condition of the patient so that the epileptor observer converges to the actual value of *z* of the patient. It is worth noting that if the discussed initial condition mismatch is small enough then the observer is shown to readily converge. The point to ponder would be whether the observer could be effectively tuned to decrease its converging time in the face of large initial condition uncertainty. This takes us to the second factor contributing to the delayed observer convergence.

**Figure 4.**
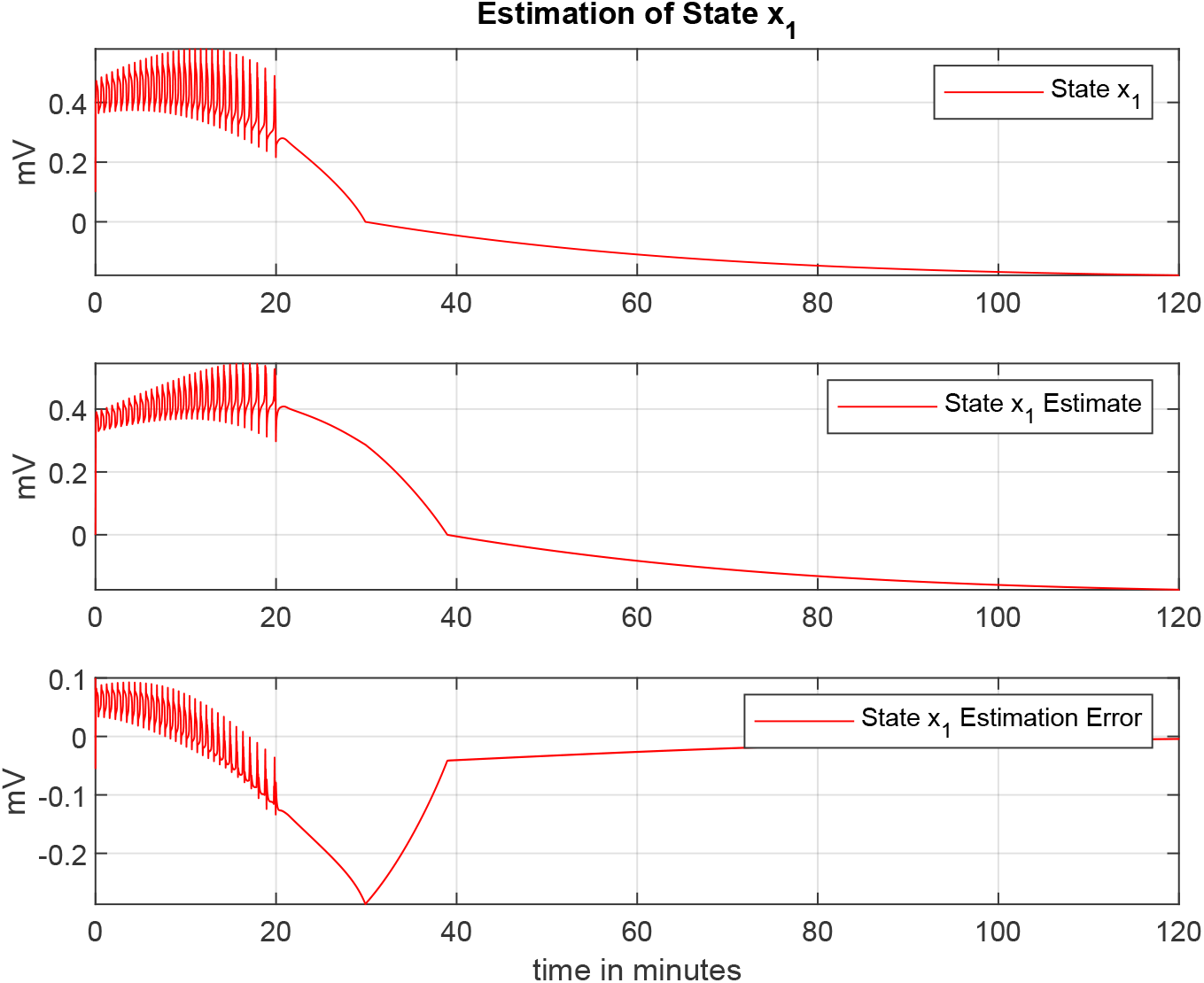
Estimation of the first state *x*_1_

**Figure 5.**
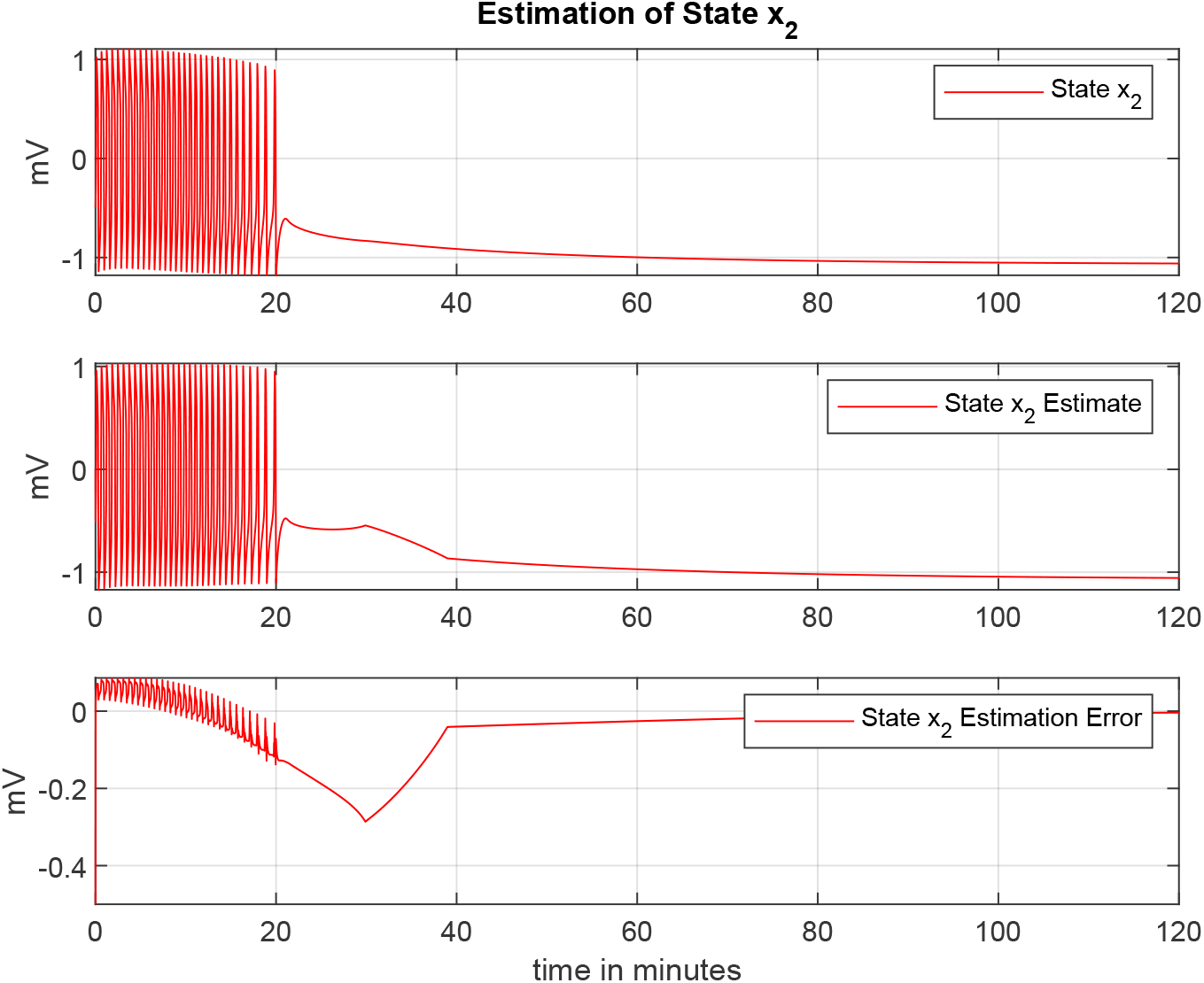
Estimation of the second state *x*_2_

**Figure 6.**
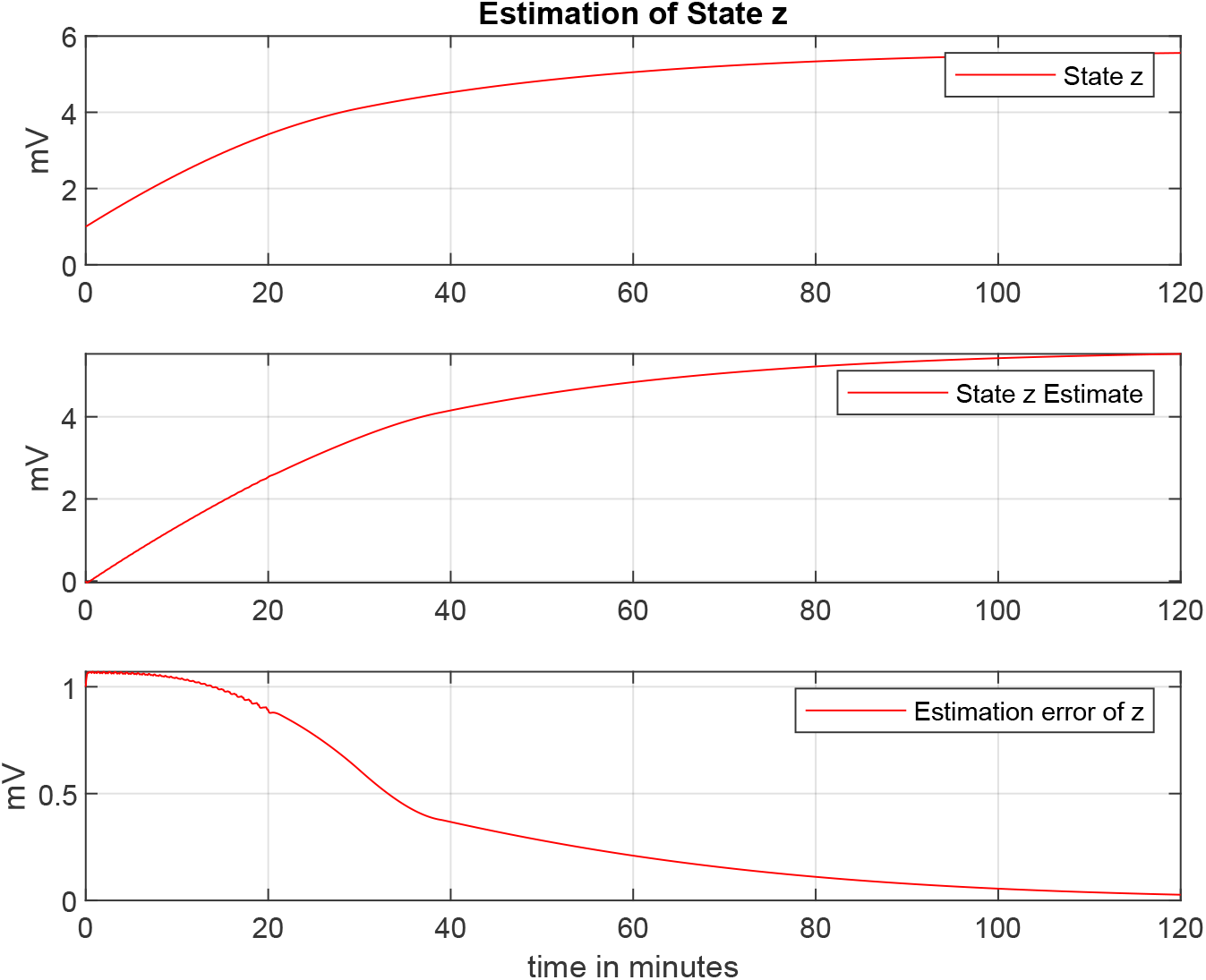
Estimation of the third state *z*

The effective tuning of this nonlinear observer loop amounts to the tuning of *R* and Lipschitz constants. While tuning, one has to take care of the condition number of the LMI solution so that it is kept robust enough against uncertainties. Secondly, there is an integrator in the epileptor model which may bar an LMI solution to have arbitrarily faster poles.

## 4. Conclusion

Despite this, it has been demonstrated that the nonlinear model of epileptor can be made the basis for designing an observer to predict the states of the epileptor. Since the epileptor represents the dynamics of the neural population of a patient whose EEG is being measured. Using the designed observer, one may obtain the synchronized epileptor which could be very useful for judging the bifurcation stage of the patient. Thus effectively predicting an upcoming seizure.

It has also been discussed that a myriad of observer types from the last eighty years of control theory have been tried on this problem during the last decade. Either these techniques depend on the linear approximation or are too complex to be implementable by COTS available computing devices or to be used by medical practitioners. It is also shown that phenomenological models like epileptor are a better bet for the time being. In the case of biological models, the real-time measurements required to determine the charge transport dynamics in a neural population are not feasible under normal clinical conditions. If the condition of neutrality is assumed in such models then these models do not remain observable as in the case of the unified neuron model proposed by Depannemaecker and his colleagues[1, 7, 8, 9].

It should be pointed out that before trying this method on the patient EEG data the unknown parameters like the time constants and the actual values of resting currents need to be estimated. Since a model structure with partially unknown parameters is at hand it would be useful to look into gray-box modeling. Pure black-box modeling methods like system identification or neural networks might fit the given EEG data but their corresponding state definitions may lack the biologically intuitive meanings.

## Notes

### Competing Interest Statement

The authors have declared no competing interest.

## References

[1] D. Depannemaecker, A. Ivanov, D. Lillo, L. Spek, C. Bernard, V. Jirsa, A unified physiological framework of transitions between seizures, sustained ictal activity, and depolarization block at the single neuron level, Journal of computational neuroscience (2022).

[2] M. Lu, Z. Guo, Z. Gao, Y. Cao, J. Fu, Multiscale brain network models and their applications in neuropsychiatric diseases, Electronics 11 (21) (2022) 3468.

[3] V. K. Jirsa, W. C. Stacey, P. P. Quilichini, A. I. Ivanov, C. Bernard, On the nature of seizure dynamics, Brain 137 (2014) 2210–2230.

[4] H. Zhang, P. Xiao, Seizure dynamics of coupled oscillators with epileptor field model, International Journal of Bifurcation and Chaos 28 (03) (2018) 1850041.

[5] K. El Houssaini, C. Bernard, V. K. Jirsa, The epileptor model: a systematic mathematical analysis linked to the dynamics of seizures, refractory status epilepticus, and depolarization block, Eneuro 7 (2) (2020).

[6] G. Ullah, S. J. Schiff, Assimilating seizure dynamics, PLoS computational biology 6 (5) (2010) e1000776.

[7] D. Depannemaecker, A. Ezzati, H. Wang, V. Jirsa, C. Bernard, From phenomenological to biophysical models of seizures, Neurobiology of Disease (2023) 106131.

[8] D. Depannemaecker, A. Destexhe, V. Jirsa, C. Bernard, Modeling seizures: From single neurons to networks, Seizure 90 (2021) 4–8.

[9] D. Depannemaecker, M. Carlu, A. Destexhe, Response dynamics of spiking network models to incoming seizure-like perturbation, Hal Science (2022).

[10] D. Fairhurst, I. Tyukin, H. Nijmeijer, C. van Leeuwen, Observers for canonic models of neural oscillators, Mathematical Modelling of Natural Phenomena 5 (2) (2010) 146–184.

[11] J. Hindmarsh, R. Rose, A model for rebound bursting in mammalian neurons, Philosophical Transactions of the Royal Society of London. Series B: Biological Sciences 346 (1316) (1994) 129–150.

[12] A. Astolfi, D. Karagiannis, R. Ortega, Nonlinear and adaptive control with applications, Vol. 187, Springer, 2008.

[13] R. Hermann, A. Krener, Nonlinear controllability and observability, IEEE Transactions on automatic control 22 (5) (1977) 728–740.

[14] A. Isidori, Nonlinear control systems: An introduction((book)), Berlin and New York, Springer-Verlag(Lecture Notes in Control and Information Sciences. 72 (1985).

[15] M. Anguelova, Observability and identifiability of nonlinear systems with applications in biology, Chalmers Tekniska Hogskola (Sweden), 2007.

[16] L. A. Aguirre, L. L. Portes, C. Letellier, Observability and synchronization of neuron models, Chaos: An Interdisciplinary Journal of Nonlinear Science 27 (10) (2017).

[17] A. Hodgkin, A. Huxley, Current and its application to conduction, J. Physiol 117 (1952) 500–544.

[18] G. A. Sotomayor, D. B. Grayden, D. Nesic, Observers for phenomenological models of epileptic seizures, in: 2023 45th Annual International Conference of the IEEE Engineering in Medicine & Biology Society (EMBC), IEEE, 2023, pp. 1–4.

[19] J. A. F. Brogin, J. Faber, D. D. Bueno, An efficient approach to define the input stimuli to suppress epileptic seizures described by the epileptor model, International Journal of Neural Systems 30 (11) (2020) 2050062.

[20] J. A. F. Brogin, J. Faber, D. D. Bueno, Burster reconstruction considering unmeasurable variables in the epileptor model, Neural Computation 33 (12) (2021) 3288–3333.

[21] J. A. F. Brogin, J. Faber, D. D. Bueno, Estimating the parameters of the epileptor model for epileptic seizure suppression, Neuroinformatics 20 (4) (2022) 919–941.

[22] S. Raghavan, J. K. Hedrick, Observer design for a class of nonlinear systems, International Journal of Control 59 (2) (1994) 515–528.

[23] R. Rajamani, Observers for lipschitz nonlinear systems, IEEE transactions on Automatic Control 43 (3) (1998) 397–401.

[24] G. Phanomchoeng, R. Rajamani, Observer design for lipschitz nonlinear systems using riccati equations, in: Proceedings of the 2010 American Control Conference, IEEE, 2010, pp. 6060–6065.

[25] S. Boyd, L. El Ghaoui, E. Feron, V. Balakrishnan, Linear matrix inequalities in system and control theory, Vol. 15, SIAM, 1994.

[26] J. Löfberg, Automatic robust convex programming, Optimization methods and software 27 (1) (2012) 115–129.

[27] X. Fan, M. Arcak, Observer design for systems with multivariable monotone nonlinearities, Systems & Control Letters 50 (4) (2003) 319–330.

[28] A. Zemouche, R. Rajamani, G. Phanomchoeng, B. Boulkroune, H. Rafaralahy, M. Zasadzinski, Circle criterion-based h-inf observer design for lipschitz and monotonic nonlinear systems–enhanced lmi conditions and constructive discussions, Automatica 85 (2017) 412–425.

[29] A. Zemouche, M. Boutayeb, A unified h adaptive observer synthesis method for a class of systems with both lipschitz and monotone nonlinearities, Systems & Control Letters 58 (4) (2009) 282–288.

[30] On lmi conditions to design observers for lipschitz nonlinear systems, Automatica 49 (2) (2013) 585–591. doi:10.1016/j.automatica.2012.11.029.

